# The dynamics of the improvising brain: a study of musical creativity using jazz improvisation

**DOI:** 10.1101/2020.01.29.924415

**Authors:** Patricia Alves Da Mota, Henrique M Fernandes, Eloise Stark, Joana Cabral, Ole Adrian Heggli, Nuno Sousa, Morten L Kringelbach, Peter Vuust

## Abstract

The neuroscience of jazz improvisation has shown promising results for understanding domain-specific and domain-general processes of creativity. Here, we used fMRI to measure for the first time the dynamic neural substrates of musical creativity in 16 skilled jazz pianists while they played by memory, improvised freely (*iFreely*) and by melody (*iMelody*), and during resting-state. We used the Leading Eigenvector Dynamics Analysis (LEiDA) to examine how different modes of improvisation (musical creativity) evolve over time, and which cognitive mechanisms are responsible for different stages of musical creation. Our results reveal that a substate comprising auditory, sensorimotor and posterior salience networks had a significantly higher probability of occurrence (POc) in both modes of improvisation than in resting-state and play by memory. Another substate comprising the default mode (DMN), executive control (ECN) and language networks had significantly lower POc in *iFreely* than in resting-state, with *iMelody* having a higher POc than *iFreely.* Such indicates that *iMelody,* a more constrained form of creativity involves a higher recurrence of subsystems responsible for goal-directed cognition and cognitive control processes. On the other hand, *iFreely* recruits brain networks responsible for generation of spontaneous musical creativity. Overall, this study brings new insights into the large-scale brain mechanisms supporting and promoting the complex process of creativity, specifically in the context of jazz improvisation, as well as the relevance of different improvisation modes in creativity research.

## Introduction

*“Jazz is not just music, it’s a way of life, it’s a way of being, a way of thinking.” – Nina Simone*

Listening to jazz musicians improvise is a spellbinding experience. Jazz musicians are able to spontaneously generate novel pieces of music in a short time frame, creating musical pieces which are both aesthetically and emotionally rewarding^1^. They must balance several simultaneous processes, involving generating and evaluating melodic and rhythmic sequences, coordinating their own performance with fellow musicians, and executing fine motor movements, all in real-time^2,3^. Jazz musicians have been found to show greater openness to experience and higher divergent thinking on personality assessments, even when compared to musicians who don’t practice jazz^4^. This phenomenal feat of human improvisation and creativity has been of great interest to neuroscientists who wish to understand the dynamics of the improvising brain, and more specifically the brain dynamics underlying the spontaneous creative process.

Creativity is often defined as “the act of creating something new and useful” ^5^, but novelty or unpredictability may not be enough. Boden comments instead on how “constraints and unpredictability, familiarity and surprise, are somehow combined in original thinking.” This distinction is important as creative music must also be aesthetically congruent with the physical constraints of the known musical range – it cannot be simply unpredictable or completely surprising. Martindale^6,7^ posited that individual differences in the breadth or narrowness of the internal attentional selection of conceptual representations may also relate to creativity. For instance, a broad focus upon conceptual concepts would activate more remote ‘nodes’ in memory. This is important as it suggests that the most creative people are those who can access associative mnemonic content in a broader way, thereby widening the constraints attached to their musical production and allowing for more unpredictable and surprising content, while the content is still familiar by its association.

Creativity can be measured by convergent and divergent thinking tasks ^8,9^. Convergent thinking consists of a single solution to a given problem, whereas divergent thinking is the generation of several different ideas to solve a given problem^10,11^. Some authors suggest the importance of both strategies in creative thinking ^9^. Creativity can be observed in numerous domains, such as in science, engineering, education and art^10,12^.

The neural signatures underlying creative thought have been investigated using diverse tasks such as artistic creativity (such as music improvisation and drawing) and general creativity (e.g. using divergent thinking tasks). Overall, the majority of the studies have used divergent thinking and creative problem solving, and only a few studies have used musically creative tasks to assess creative thought more generally^13,14^. The idea that improvisation involves divergent thinking processes has also been suggested ^15,16^. Pressing ^17^ proposed that improvisation results from the continuous recall of previous information (familiar music structures) and rearrangement of this information in a flexible manner in order to create something new (unfamiliar/surprising) ^17^. Thus, musical improvisation involves process-specific divergent thinking. The neuroscience of musical creativity has thus far shown promising results for understanding not only domain-specific creative thought, but also domain-general processes of creativity^1–3,18,19^.

Jazz improvisation is well-suited for studying spontaneous creativity due to its reliance upon known neural and cognitive processes. Jazz improvisers spontaneously evaluate and generate melodic, rhythmic and harmonic patterns, enacted with fine motor movements, in order to create novel musical sequences in real time ^2,3,20,21^. Jazz musicians have shown to triumph in domain-general creative abilities ^4,22,23^, to have a greater sensitivity to unpredictable sound changes ^23,24^, and a higher intrinsic motivation, and pleasure in musical activities ^25^, even when comparing with other musicians.

The growing interest in the neuroscience of creativity has led researchers to explore the brain structure and dynamics of creative thinking, and despite the heterogeneity of neuroimaging studies in domain-general and artistic creativity, the patterns of brain activity and connectivity reported are similar. Network neuroscience defines the brain as a graph or network ^26^, where “edges” are used to represent structural or functional connections between brain regions (known as “nodes”). Creativity studies have also started to uncover the brain networks underlying the complex creative process in divergent thinking ^27^, as well as connectivity patterns in resting-state fMRI in creative individuals ^28,29^. Several studies using divergent thinking tasks in functional magnetic resonance imaging (fMRI) have found creativity to be the result of a dynamic interplay between different brain networks ^2,19,30–32^, namely the default mode (DMN), executive control (ECN) and salience network (SN) ^28,33,34^.

The DMN is known to be linked to spontaneous and self-generated cognition, such as in daydreaming, mind-wandering, episodic memory retrieval and future thinking ^13,28,35,36^, and DMN activity during the creative process has been suggested to reflect the spontaneous generation of ideas, acquired with the aid of long-term memory ^28^. The ECN is linked to goal-directed cognition and cognitive control processes such as working memory and response inhibition ^37^ playing a key role in creativity through guiding, constraining, and modifying DMN processes in order to meet the individual’s creative task goals ^38–41^. The SN is generally known to facilitate coupling between default and control networks, thus being able to support the dynamic interplay between evaluative and generative processes during the creative process ^42^.

Although there is a consensus about the involvement of these networks in creativity, their role in musical creativity ^3,21,32,43–47^ is still unclear and the literature presents inconsistent results. While some studies showed increased activity in the DMN ^32,47^, others have reported the opposite^43,44^. The same has been documented for the ECN, more specifically for the dorsal lateral prefrontal cortex (DLPFC)^32,48^. Other brain networks (regions) which are also found to be involved in creative thought have been associated with different cognitive processes, such as attention and executive control, motor sequence generation, voluntary selection, sensorimotor integration, multimodal sensation, emotional processing and interpersonal communication^19,31,49^. Nonetheless, as Bashwiner and colleagues ^50,51^ outlined, it is surprising that in studies of musical creativity, auditory networks which are expected to have a strong link with musical creativity are not found to be significantly more active when compared to a control task.

Another important issue concerns the strategies that jazz musicians use for improvisation. The most common strategy is to improvise *freely* but according to a chord scheme belonging to the specific tune they are playing^24^. Here, jazz musicians use their skills and previously practiced melodic and harmonic material as building blocks with the aim of creating musical pieces that are novel and engaging^52,53^. Consequently, this approach may entail brain processes similar to the ones underlying divergent thinking^2^. Another oft used strategy is to use the melody as the starting point for the improvisation^53^. Here the outcome usually becomes less complex, and more ‘hummable’ and may as such be more related to emotional processing which is known to be associated with the perception of songs. Many jazz musicians who are proficiently using this approach are known to accompanying their instrumental improvisation with vocalization (such as Keith Jarrett)^54^. Since this approach involves a goal-oriented task it may be closer related to convergent thinking than free improvisation.

Previous studies have shown that creativity is a result of a dynamic interplay between different brain networks ^2,30,32^, however none have yet explored the brain functional dynamics of spontaneous musical creativity through jazz improvisation. Here we propose to explore, for the first time, the whole-brain dynamics underlying spontaneous musical creation in jazz pianists, using Leading Eigenvector Dynamics Analysis (LEiDA)^55–58^. LEiDA captures the instantaneous BOLD phase signal and uses leading eigenvector decomposition to find the recurrent functional connectivity patterns (or brain substates). In this study, we quantified the differences in terms of probability of occurrence, using two complementary tasks during fMRI (two different modes of musical improvisation: one constrained by melody and one freely), and we also measured a play by heart (memory condition) and resting state during the same MRI session (rs-fMRI) as a baseline.

We hypothesised that given how musical creativity is a rich and complex dynamic process, we would find corresponding signatures of brain dynamics (recurrent Phase Locking – PL - metastable substates) that are significantly altered when compared to a baseline condition of playing (memory) and resting state. We further hypothesised, based on Loui’s ^19^ model of musical creativity, that different connectivity patterns would be associated with distinct stages of music creation – idea generation, perception and evaluation – during improvisation, and that these connectivity patterns would be different when improvising freely (*iFreely*), which has a higher level of freedom, than when improvising constrained by the melody *(iMelody).*

## Results

In this study, we investigated the dynamic nature of the jazz musician’s brain while they improvised by melody and freely, by characterising the most recurrent patterns of whole-brain functional connectivity arising during the six minutes of playing in each condition.

### Detection of the PL substates

The repertoire of metastable substates depends upon the number of clusters determined by the k-means clustering algorithm, where higher number of clusters usually results in less frequent and more fine-grained substates ^58^. In this study, we did not aim to determine the optimal number of PL substates but rather to search for the PL substates, which significantly and recurrently characterize musical improvisation, using a memory condition and resting-state as a baseline.

To do this, we first estimated the PL substates that most significantly and consistently differentiated improvisation (both modes of improvisation) from the baseline condition (resting-state). Across all 13 partition models produced, a total of 12 PL substates were found to be significantly different (Bonferroni corrected) between improvisation modes and resting-state in terms of probability of occurrence. For each partition model, statistical significance between pairs of conditions (in terms of probability and duration of the PL substates) was assessed using permutation-based paired t-tests between-condition comparison (Figure 1 left), and corrected for multiple comparisons using Bonferroni’s method). Based on the criteria of the minimum number of PL substates revealing significant differences in POc between all music-playing modes under study and the baseline condition, we selected the partition solution that divides the brain dynamics of all conditions into five PL substates (k=5). In other words, the BOLD PL patterns captured for all time-points, conditions, and subjects are clustered into five cluster centroids. The partition into five substates is in line with the literature, where 5 to 10 functional networks emerge during rest ^55,59^. Furthermore, as highlighted in Figure 1 (left), all five substates derived at the chosen k-level consistently reappeared across multiple partition solutions (DSC ≥ 0.8; k ∈ [5 .. 9]), thus strengthening the reliability of this method and criteria used in capturing a robust signature of brain dynamics across all conditions under study.

**Figure 1.**
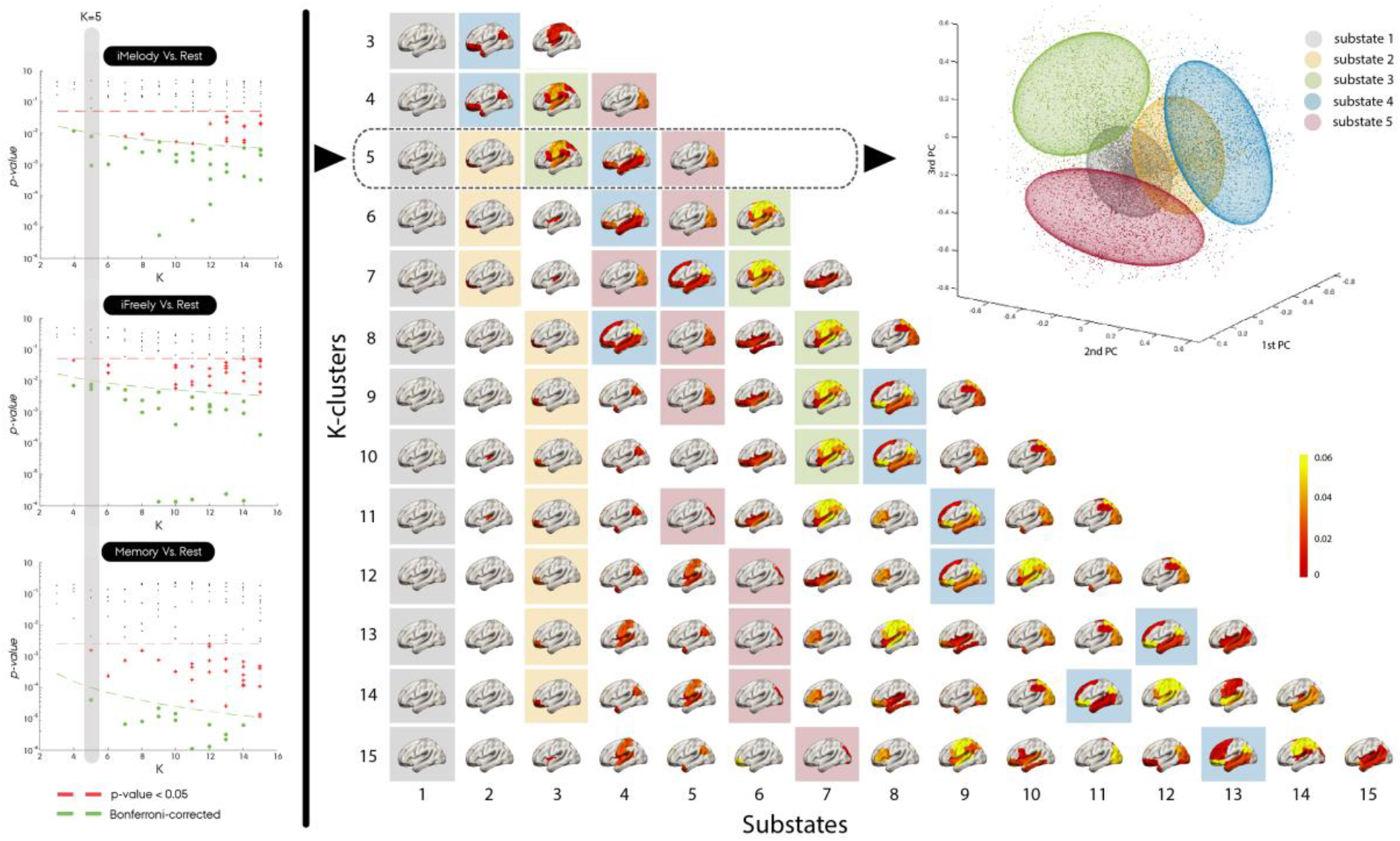
Detection of significant between-condition differences in phase-locking (PL) substates across the 13 partition models. Left) For each pair of conditions and partition model (k-means clustering solution), we plot the p-values associated with the between-condition comparison between experimental condition and rest in terms of probability. The p-values marked as green dots survive the correction for multiple comparisons within each partition model (<0.05/k). The grey-shaded column highlights the partition model chosen (k=5; the lowest k-value with significant statistical difference between each of the music playing conditions and the rest condition) to describe the clustered dynamics of the experimental conditions under study. Middle) Brain rendering of the PL substates across partition models. Brain regions are colour according to PL amplitude of the eigenvector representing them. Here, we used the Dice Similarity Coefficient (DSC) to highlight (background square) how consistent the substates are across k-solutions, with regards to k=5. Using the reference substate colour-coding of k=5, across clustering solutions, substates are highlighted with the corresponding colour of the reference substate if it shares a DSC ≥ 0.8. Right) The BOLD PL patterns (leading eigenvectors of BOLD phases) captured for all time-points, conditions and subjects can be represented in a three-dimensional version of the phase space. Here, each data point (dot) is placed according to their cosine distance to the three principal components, or eigenvectors of the covariance matrix, estimated from all observations. Observations are coloured according to the cluster they are assigned to for k=5 (i.e. the closest cluster centroid). Colour-coding is congruent with the brain renderings projecting each of the cluster centroids (i.e. each PL substate). Additionally, ellipsoids are fitted to each set of data points to represent the degree of dispersion and directionality of each cluster cloud.

### Repertoire of recurrent PL substates

In line with previous studies using LEiDA ^55–58^, the most probable state of BOLD phase-locking is a global substate, where all BOLD signal are synchronized. The remaining four recurrent substates were found to overlap with typical RSNs reported in the literature ^60,61^.

### Probabilities of occurrence – what is special about improvisation?

We found a recurrent substate, substate 3, with significantly higher probability of occurrence for both modes of improvisation (*iMelody* and *iFreely*) compared to rest (Figure 2). This PL substate includes the bilateral: precentral gyrus, Rolandic operculum, posterior cingulate gyrus, superior parietal, supramarginal gyrus, Heschl’s gyrus, superior temporal gyrus; the left: inferior frontal gyrus opercularis, paracentral lobule and pallidum; and the right insula. These brain regions are mainly part of the auditory, sensorimotor and the posterior salience networks.

**Figure 2.**
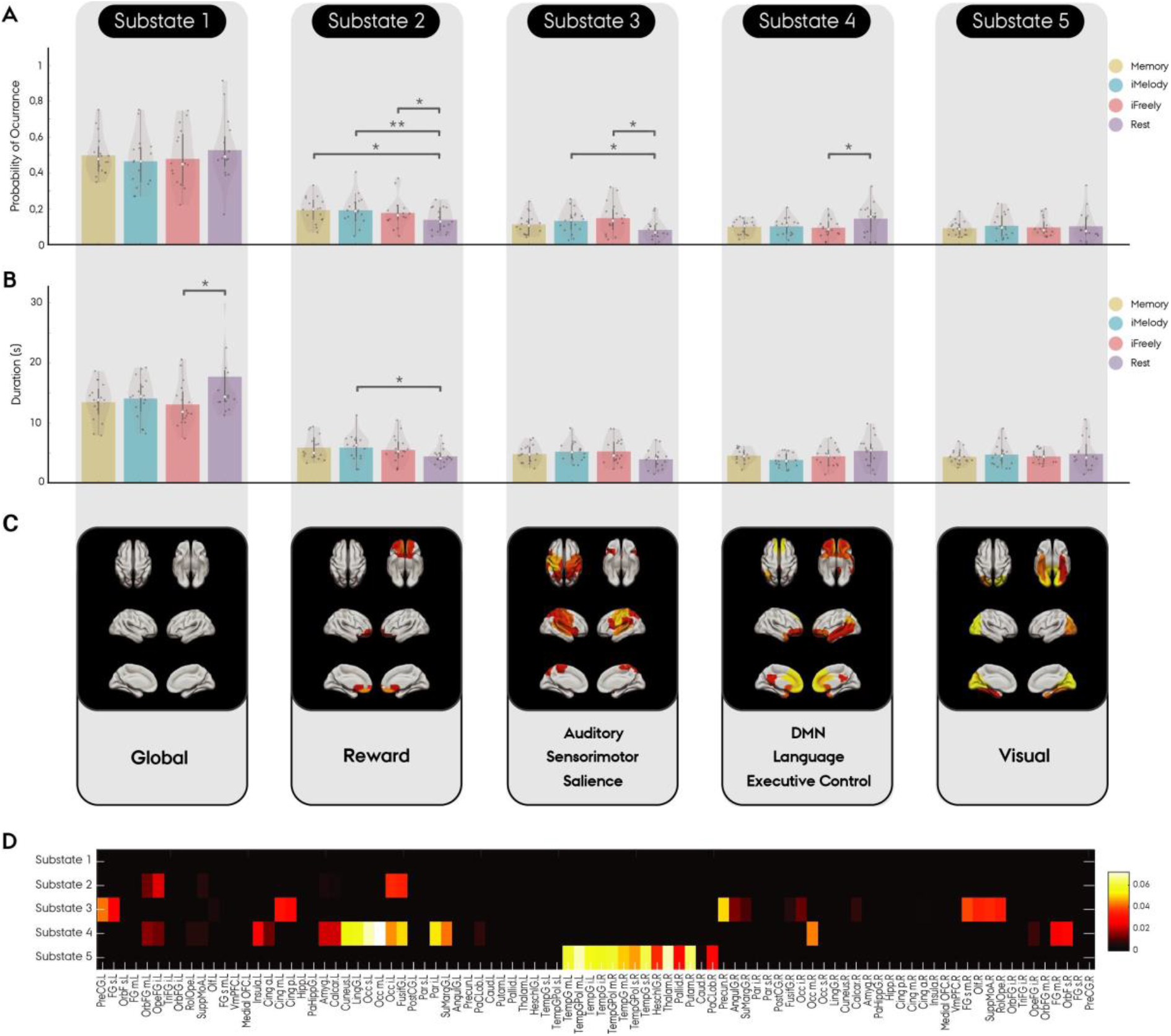
Signature of Musical Creativity. Characterising the repertoire of metastable substates during jazz improvisation, play by memory and resting-state. **A)** Probability of occurrence (POc) of each of the five brain substates estimated using LEiDA, during play by memory (yellow) improvisation on the melody (cyan), improvisation freely (red) and rest (grey), represented by bar and violin plots. Substate 3 was found to have significantly (p<0.05; Bonferroni-corrected) higher POc in both modes of improvisation (iMelody and iFreely) than in resting-state and Substate 4 had significantly lower POc in iFreely than in resting-state. **B)** duration of each of the five brain substates. **C)** Rendering of the brain, showed as top, bottom (whole) and side (hesmispheric) planes, for the five substates and corresponding RSNs. **D)** participation and connection weight of AAL regions in each of the five substates. Our analysis revealed five recurrent PL substates, one global (substate 1) and four recurrent substates, reflecting: reward (substate 2), an auditory-motor network (substate 3), a complex array of functions that support improvisation and creativity more generally (such as evaluation-perception) (substate 4) and visual imagery (substate 5).

Interestingly, another recurrent substate, substate 4, was found to have lower probability of occurrence (but with higher duration) in *iFreely* than in *iMelody,* and significantly lower probability of occurrence in *iFreely* than in rest. This PL substate includes the bilateral: orbital frontal cortex (inferior, middle and superior), olfactory cortex, medial prefrontal cortex, ventromedial prefrontal cortex, medial orbitofrontal cortex, anterior cingulum, and middle temporal gyrus; the left: posterior cingulum, angular gyrus and the superior temporal gyrus. These brain regions are mostly part of the dorsal default mode (DMN), language and the left executive control (ECN) networks.

## Discussion

The present study investigated the whole-brain dynamics underlying musical improvisation in a group of jazz pianists. A novel analytic method – LEiDA – was used to estimate the metastable brain substates (i.e. recurrent patterns of brain functional connectivity) in which the brain is characterized during two types of musical improvisation: freely and by melody, playing by heart the melody from “The Days of Wine and Roses” (memory condition), as well as at rest. Our dynamic analyses show a recurrent substate – “improvisation mode” – characterised by a significantly higher probability of occurrence in both modes of improvisation (*iMelody* and *iFreely*) when compared to playing by memory and resting-state. This substate comprises mainly areas of the auditory, sensorimotor and posterior salience networks. Our analyses also revealed a recurrent substate comprising mainly areas of the dorsal default mode (DMN), the left executive control (ECN) and the anterior salience (SN) networks with significantly higher probability of occurrence in *resting-state* than in *iFreely* (a more free mode of improvisation)*, with iFreely* having the lowest probability of occurrence of all the four conditions.

In sum, jazz improvisation relies strongly on the use of multiple brain substates (each, represented by a network combining multiple fundamental brain mechanisms), recursively, when they improvise in order to create music, which is novel, surprising, aesthetically balanced and emotionally rewarding. Moreover, we provide the first comprehensive characterization of the functional brain networks involved in musical creativity, as well as the changes in their dynamic fingerprint (i.e. probability of occurrence) compared to two baseline conditions (playing by memory and during the resting state).

Here, we describe our findings in terms of the differences in dynamic structure, by first characterizing the five brain substates that best describe, musical improvisation, play by memory and resting-state in jazz musicians, and second, by exploring the probability of occurrence of these substates.

We selected the model reflecting five functional substates, as it revealed significant differences in the probability of occurrence between the four conditions of interest. Our partition model of five substates is in accordance with the literature, where 5 to 10 functional networks emerge during rest ^55,59^. Our results revealed, in line with previous studies using LEiDA ^55–58^, a global substate (substate 1), but also other four recurrent substates which overlap with resting-state networks (RSNs) reported in the literature ^60,61^. These four recurrent substates reflect: reward (substate 2), an auditory-motor network in substate 3, a complex array of functions that support music creation on the “fly”, involving intuitive decision making (evaluation/perception) in substate 4, and a visual network – visual imagery – in substate 5.

Our dynamic analysis revealed that substate 3 had a significantly higher probability of occurrence, for both modes of improvisation (*iMelody* and *iFreely*) when compared to resting-state. This substate comprises brain regions including the bilateral: rolandic operculum, pre- and post-central, superior parietal, supramarginal, heschl and superior temporal gyrus; the left: paracentral lobule, pallidum and inferior frontal gyrus opercularis; and the right insula. This substate covers mainly areas of the auditory, sensorimotor and posterior salience networks.

Sensorimotor regions have been reported to be involved in artistic creativity studies. A recent meta-analysis in artistic creativity (music, literary and drawing) reported common domain-general brain activity patterns in the pre-supplementary motor area, left dorsolateral prefrontal cortex and right inferior frontal gyrus (IFG) across the different types of creativity^62^. However, the authors also found domain-specificity musical creativity activity in the left: supplementary motor area (SMA), precentral gyrus and middle frontal gyrus, and bilateral inferior frontal gyrus^62^. In musical creativity, an increased connectivity of sensorimotor regions have been linked to higher levels of expertise in improvisation, when comparing musicians with different levels of expertise in improvisation^43^. It is known that sensorimotor networks play an important role in music improvisation, however it is surprising that in studies of musical improvisation, auditory networks are not found to be significantly active when compared to a control task^50^. Bashwiner and Bacon ^51^ reported that in 16 musical creativity studies all have implicated motor regions, but only half of them have reported auditory-associative regions to be implicated in musical creativity. This is despite jazz musicians reporting that they “hear” (auditory imagery) what they intend to play even before playing it, i.e. form predictions about what a piece of music will sound like even before the actual performance. Thus, auditory networks are expected to have a strong link with musical creativity. Motor imagery may also be expected, as musicians may simultaneously imagine themselves performing the motor sequence involved in playing their chosen instrument ^21,47,52,64^.

In addition, we found differences in the probability of occurrence when improvising by melody or improvising freely. Melodic improvisation (*iMelody*) involves a goal-specific task of arranging the notes in a certain order trying to create a new melody that bears resemblance to the original, in this case a known musical song (DWR). On the other hand, the free improvisation (*iFreely*) in this study comprises a chord scheme, allowing for the use of a larger repertoire of melodic and rhythmic material, and preliminary analysis (to be published elsewhere) revealed that participants played a higher number of notes in *iFreely* than in the *iMelody* condition. In this study, the *iFreely* condition corresponds closely to the unconstrained improvisation performed by jazz musicians in a natural playing situation, whereas the *iMelody* leads to a more constrained improvisation.

Another recurrent substate – substate 4 – with significantly lower probability of occurrence for *iFreely* than for the resting-state (with resting-state having the highest probability of occurrence for all four conditions) was also found. This substate comprises brain regions of the dorsal default mode (DMN), language and the left executive control (ECN) networks, including the bilateral: ventromedial prefrontal cortex, medial prefrontal cortex, all sections of the orbital frontal cortex, olfactory cortex, middle temporal poles, anterior cingulum; and the left: angular gyrus, posterior cingulum and the middle temporal gyrus. These results are in line with previous research, where the DMN (responsible for spontaneous and self-generated thought) and the ECN (responsible for cognitive control and maintenance of goal-directed cognitive processes) are shown to cooperate in order to generate and evaluate ideas during creative tasks ^2,30,43^.

In sum, the regions belonging to substate 4, which has a significantly higher probability of occurrence in resting-state than in improvisation, describes an interaction between different cognitive processes such as idea generation (where attention, memory retrieval and mind-wandering are needed), selection, evaluation (of idea and reward value) and perception. In music improvisation this lower predominance may be explained by the fact that in order to overcome knowledge constraints a down-regulation of prefrontal cortex regions (reduced cognitive control) and lower recurrence to salient information is needed ^10,65^. Lesion studies in the dorsolateral PFC and orbitofrontal cortex (OFC) have been related to higher performance on problem-solving tasks^66^ and to an increased capacity to overcome salient information during the idea generation stage^65^. Of note, this substate was found to have a slightly higher dominance in *iMelody* than in *iFreely*, which suggests that different improvisational strategies may rely upon different cognitive processes.

Our results also revealed a substate comprising limbic regions, such as the bilateral superior, medial, and middle orbitofrontal and olfactory cortices. These regions cluster bilaterally around the orbitofrontal cortex, a region known to be a nexus for sensory integration, prediction-monitoring, and reward^67^. This could reflect the need for the improvised music to remain emotionally compatible with the preceding musical elements. A further possibility is that this activity reflects the musician’s emotional experience and investment in the process of improvisation. However, while some studies note significance of limbic regions in musical improvisation ^19,68^, there are no reports of these regions being significantly active in studies of musical creativity. This may be explained by the fact that researchers usually use another playing condition to contrast with an improvisation task in music creativity studies^2,68^. Here, the connectivity of substate 2 therefore leaves it in an important position for integrating the sensory features of incoming musical stimuli, and the rewarding elements of playing music (either by memory or improvisation) – both monitoring and predicting the reward value of musical features, and subjective enjoyment of playing music. This substate was found to have significantly higher probability of occurrence in all playing conditions compared to the resting-state, with the playing by memory condition having the highest probability of occurrence. Music is known to be as rewarding as food or sex ^69^, and is rated to be on the top 10 things that people find most pleasurable in life^70^. Creating something that has an impact in people’s life, such as music, is of great importance for musicians, who have a higher intrinsic motivation and pleasure in musical activities, particularly jazz musicians ^24^. This opens up to the question if reward, pleasure, and the drive to create are linked and essential for the creative process to happen or if it is a domain-specific process of musical creativity.

Interestingly, another substate comprising brain regions mainly belonging to the visual network, such as the bilateral: calcarine, cuneus, lingual, occipital (inferior, middle and superior) and the fusiform gyrus was found to have similar probabilities of occurrence in the four different experimental conditions. Areas in the occipital cortex (visual network) have been found active in diverse studies of jazz improvisation ^3,20,32^. One case-study used a classical musician with great improvisational skills suggested that the activation of the visual cortex could be related to visual imagery (i.e. the capacity to visualize image representations in the mind in the absence of physical stimuli), but the author speculated that this was due to the participant being able to see her hands during the performance ^71^. In our study the participants were not able to see their hands during the performance, suggestive of a different role of visual imagery during improvisation. Anecdotal reports from a diverse sample of creative individuals points to the importance of such imagery within the creative process. These reports have been made in different creative domains, such as music, science, art, and literature ^72,73^. Visual imagery may be of high importance during the creative process as musicians recurrently listen to and create music mentally ^71,74^. It may also support the divergent exploration and generation of new musical ideas, thereby facilitating the psychological evaluation of the combination of familiar musical structures in an unfamiliar structure, leading to the creative output. Visual imagery ability has also been shown to be of benefit in domain-general creative tasks, such as divergent thinking ^73^.

A limitation of this study is that we can but make inferences about domain-general creativity. Future studies will need to compare improvisation in different modalities – perhaps verbal (divergent thinking), auditory (music), visual (art), and kinaesthetic (dance) to confirm whether the network herein does indeed support domain-general creativity. For future work in this area, we would suggest that extending this novel approach, LEiDA, to parcellation schemes with a higher number of areas than that supported by the Automated Anatomical Labelling (AAL) atlas. For example, the parcellation schemes of Shen and colleagues’ ^75^ or Glasser and colleagues’ ^76^, could uncover more fine-grained substates.

In summary, this study provides a novel approach to studying the brain dynamics of musical creativity. Jazz improvisation reflects a complex and multifaceted set of cognitive processes that have correspondingly complex functional network dynamics. Here, we attempted for the first time to unravel the dynamic neural cognitive processes involved in musical improvisation over time. We show that different cognitive processes are involved in musical creativity and supported by different brain networks, including the motivation to create (limbic network), divergent thinking processes responsible for mental exploration and generation of new combinations of familiar musical sequences (auditory-motor), convergent thinking processes to evaluate and revise the predicted initial musical thoughts to create an aesthetically musical improvisation (DMN-ECN) and visual imagery responsible for visualizing new combinations of musical sequences (visual network) are involved in musical creativity. These five brain substates also support the model of improvisation proposed by Pressing, where improvisation is described to be a dynamic interplay between motivation, generation, evaluation and execution of novel motor sequences ^17^. In addition, we have also shown that musical creativity is facilitated by interactions between convergent and divergent thinking, where musicians need to explore multiple musical possibilities, but with a goal of arriving at a musical output that is novel and aesthetically pleasant. We found that improvising on the melody and improvising freely on a harmonic progression shared a common fingerprint of brain substates underlying the process of musical creation characterized by the brain spending more time in a substate linked to the functional role of auditory-sensorimotor and reward networks. This may reflect functions such as generating and evaluating creative ideas, predicting and monitoring sensory input, and syntactic processing through linguistic mechanisms. In comparison to melodic improvisation (resting-state and play by memory conditions) the act of improvising more freely was characterized by a lower predominance (recurrence) of default mode, language and executive control networks. These results show the benefit of using novel methods (such as dynamic functional connectivity) and musical paradigms to investigate the large-scale brain mechanisms involved in the complex process of musical creativity, and creativity in general.

## Methods

### Participants

The total sample consisted of 24 right-handed male musicians with normal hearing and no history of neurological disease. Eight participants were excluded from the analyses: 2 found out that they were claustrophobes and 6 were excluded due to excessive head movement. Our final sample resulted in 16 participants (mean 28.0 ± 8.71 SD). All participants were proficient in jazz piano playing (with at least 5 years of experience), and they stated that they practice on average 1.9 ± 0.9 SD hours per day, and 22 ± 7.7 days of practicing per month. All participants gave written consent to participate in the study. The study was approved by the local ethics committee and it was undertaken in accordance with the Helsinki declaration.

### Stimuli and procedure

We acquired functional MRI while participants were playing on an MRI compatible keyboard in four different conditions in a pre-defined randomized order, while listening to the chords of the jazz standard “The days of wine and roses” (DWR). Participants were asked to:

a. play the melody of DWR by memory (Memory)
b. play from a score sheet (Read) which was an alternative melody composed specifically for this experiment on the chord scheme of DWR
c. improvise on the melody (*iMelody),* i.e. play melodically as if they were to create a new melody for the chord scheme of DWR
d. improvise freely on the chord scheme for DWR (*iFreely*).

Each condition lasted for 45 seconds, and participants had to play it eight times. In total each participant played 24 minutes (6 minutes for each condition) (Figure 3-A).

**Figure 3.**
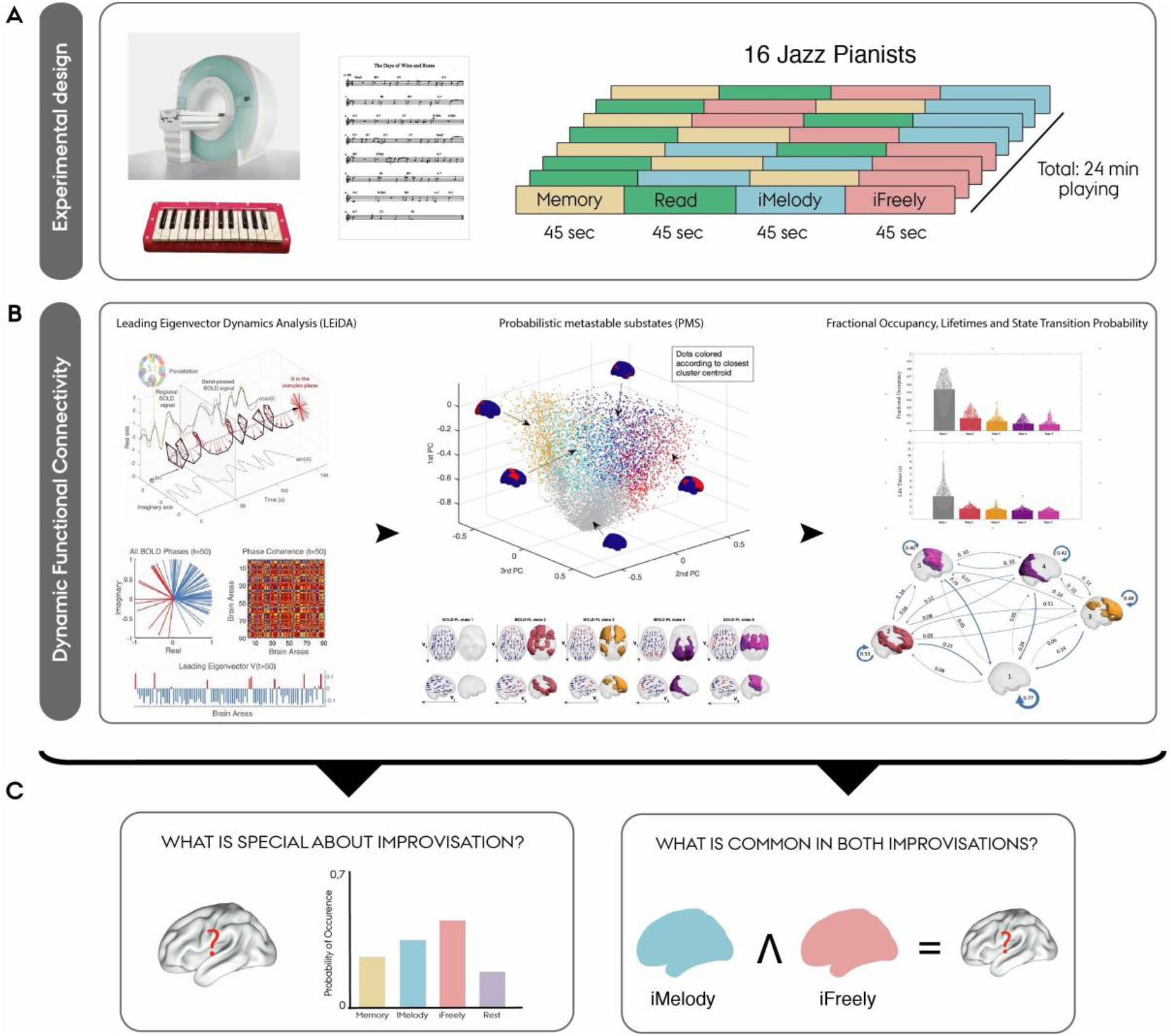
The dynamics of the improvising brain: experimental protocol and methods. A) Experimental design: participants were asked to play four different conditions inside of the MRI scanner using a 25 keys MRI-compatible keyboard. The four different conditions were: play by memory (Memory), play from a score sheet (Read), improvise by melody (iMelody) and freely improvise (iFreely). B) LEiDA (Leading eigenvector dynamics analysis) captures the BOLD phase locking (PL) of the system focusing on the dominant pattern captured by the leading eigenvector of dynamic PL matrices. C) The aims of the current study were centred in providing a better understanding of the fingerprints of brain dynamics that are specific to the process of improvisation, as well D) which features were common or exclusive to different modes of improvisation.

To ensure that there were no image artefacts, we used a custom-made MR-compatible fiber optic piano keyboard ^77^. The keyboard, consisting of 25 full size keys, covered two full octaves, and its lightweight and slim design allowed it to be positioned on the participants’ laps, such that all keys could be reached by moving only the forearm. Participants were instructed to only play with their right hand. Output from the keyboard was interpreted into a MIDI signal by a microcontroller outside of the scanner room. Piano sounds were generated by a Roland JV-1010 hardware synthesizer based on this MIDI signal. The piano sound from the synthesizer was subsequently mixed together with a backing track, and delivered to the participants through OptoACTIVE noise cancelling headphones.

The instructions for each condition were controlled by a PsychoPy ^78^ script on a laptop computer. A MR compatible screen was used to project the instructions and participants viewed it using a mirror that was attached to the head coil. Participants were instructed about the conditions before going inside of the scanner, and they were allowed to play 2 times the score sheet outside the scanner, to make sure they would understand that they needed to read from a score inside the MR scanner. Inside the scanner participants received the information about which condition they should play through the screen.

### Image acquisition and processing

All participants underwent the same imaging protocol using a 32-channel head coil in a Siemens 3 T Trim Trio magnetic resonance scanner located at Aarhus University Hospital, Denmark. Whole-brain T1-weigthed and task-based fMRI images were acquired for each participant.

### Anatomical scan acquisition

The 3D T1-weigthed sequence was performed with the following parameters: sagittal orientation; 256 x 256 reconstructed matrix; 176 slices; slice thickness of 1 mm; echo time (TE) of 3.7 ms; repetition time (TR) of 2420 ms; flip-angle (α) of 9.

### fMRI Acquisition

A multi-echo EPI-sequence was acquired with a total of 1484 volumes and with the following parameters: voxel size of 252 x 252 x 250 mm; 54 slices; slice thickness of 2.50 mm; multi-echo time: TE1= 12 ms, TE2= 27.52 ms, TE3= 43.04 ms, TE4= 58.56 ms; repetition time (TR) of 1460 ms; flip-angle (α) of 71. Only the second echo was used in our analysis.

### fMRI Processing

The fMRI data was processed using MELODIC (Multivariate Exploratory Linear Decomposition into Independent Components)^79^ part of FSL (FMRIB’s Software Library, www.fmri.ox.ac.uk/fsl). The default parameters of this imaging pre-processing pipeline were used for all the 16 participants: motion correction using MCFLIRT ^80^; non-brain removal using BET ^81^; spatial smoothing using a Gaussian kernel of FWHM 5 mm; grand-mean intensity normalization of the entire 4D dataset by a single multiplicative factor and high pass temporal filtering (Gaussian-weighted least-squares straight line fitting with sigma = 50 seconds). FSL tools were used to extract and average the time courses from all voxels within each cluster in the AAL-90 atlas ^82^.

### Dynamic Functional Connectivity Analysis

We applied a data driven approach to capture BOLD phase-locking (PL) patterns from fMRI data at single TR resolution with reduced dimensionality, the Leading Eigenvector Dynamics Analysis (LEiDA)^55^. On a first stage, the BOLD signals in the *N*=90 brain areas were band-pass filtered between 0.02 Hz and 0.1 Hz and subsequently the phase of the filtered BOLD signals was estimated using the Hilbert transform ^55,83^. The Hilbert transform expresses a given signal x as x(t) = A(t)*cos(θ(t)), where A is the time-varying amplitude and θ is the time-varying phase (see Figure 3-B left). We computed the dynamic phase-locking matrix (dPL, with size NxNxT). Each entry of the dPL(n,p,t) estimates the phase alignment (which varies from −1 to 1) between brain regions n and p at time t, given by the following equation:

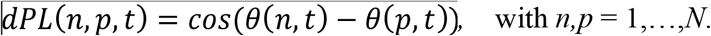

To characterize the evolution of the dPL matrix over time with reduced dimensionality, we considered only its leading eigenvector, *V1(t*), which is a *Nx1* vector that captures, at time *t,* the projection of the BOLD phase in each brain area into the main *orientation* of BOLD phases over all areas (Figure 3-B, second panel from the left). When all elements of *V1(t*) have the same sign, all BOLD phases project in the same direction with respect to the orientation determined by *V1(t).* If instead the first eigenvector *V1(t*) has elements of different signs (i.e., positive and negative), the BOLD signals project into different directions with respect to the leading eigenvector, which naturally divides the brain into distinct modes (colored in red and blue in Figure 3-B second panel from the left). Previous studies using LEiDA have shown that the subset of brain areas whose BOLD signals appear temporally phase-shifted from the main BOLD signal orientation reveal meaningful functional brain networks ^55–58^.

### Recurrent BOLD PL Substates

In this work, we aimed to investigate the existence of specific patterns of BOLD PL substates, or PL substates, associated with musical creativity in jazz improvisation. To do so, we first searched for recurrent BOLD phase-locking patterns emerging in each of the three experimental conditions (*Memory*, *iMelody* and *iFreely*), and compared their probabilities of occurrence to a common resting-state baseline. Recurrent BOLD PL patterns, or PL substates, were detected by applying a k-means clustering algorithm to the set of leading eigenvectors, *V1(t),* associated to the fMRI volumes acquired during each condition over all participants (considering a lag of three TR to account for the delay in the hemodynamic response), as well as the fMRI volumes recorded during a baseline (resting-state) period of 542 seconds (the same baseline was used for all 3 experimental conditions). The k-means algorithm clusters the data into an optimal set of *k* clusters, where each cluster can be interpreted as a recurrent PL substate.

While resting-state fMRI studies have revealed the existence of a reduced set of approximately 5 to 10 functional networks that recurrently and consistently emerge during rest across participants and recording sites ^55,59,61,84^, the number of PL substates emerging in brain activity during a task is undetermined, and depends on the level of precision allowed by the spatial and temporal scales of the recordings. In the current study, we did not aim to determine the optimal number of recurrent PL substates detected in a given condition, but instead to detect which clusters of the PL substates whose probability of occurrence was significantly (Bonferroni correction adjusted p-value < 0.05) and consistently modified by the experimental conditions with respect to the baseline. In that direction, we ran the k-means algorithm for 13 partition models by varying the number of clusters *k* from 3 to 15, with higher k resulting in more fine-grained configurations. Particularly, for each partition model, clustering the 239281329 leading eigenvectors, V1, (resulting from 16 participants and X TRs each for each of the four experimental conditions) returns *k* N X 1 cluster centroids, Vc, each representing a recurrent substate of BOLD PL. The brain substate related to each Vc, can be represented as a network in a cortical space. Previous works ^56,85^ have demonstrated that the smallest synchronized community in Vc has a highly significant overlap with functional networks.

### Probability of Occurrence (POc)

Recurrent substates were compared in terms of their probabilities of occurrence (calculated as the number of time points in which a PL substate in active during the scan, divided by the total number of time points in a scan) in both modes of improvisation (by melody and freely) and play by memory (Memory) with respect to their probabilities of occurrence during the condition and resting-state baseline, using a permutation-based paired t-test to assess the statistical differences. The significant thresholds were corrected to account for multiple comparisons as *0.05/k,* where *k* is the number of substates (or independent hypothesis) tested in each partition model ^56–58^.

### Comparison with resting-state networks

We used the large-scale resting-state networks (RSNs) described by Shirer and colleagues ^60^ to quantify the representation of each RSN in each of the five substates. Intersection of each of the 14 RSNs with the 90 AAL brain regions was computed. Quantification of each RSNs representation was then calculated dividing the results of the intersection between RSNs and 90 AAL by the total number of voxels of each RSNs intersected with the 90 AAL regions.

## Notes

### Competing Interest Statement

The authors have declared no competing interest.

